# Native oligomerisation of a type I antifreeze peptide from winter flounder

**DOI:** 10.64898/2026.09.25.754443

**Authors:** Lara L. Virgilio, Avend Omar, Xing Jian Chang, Jordan Stolow, Min Lee, Abiaz Hossain, Bethany Kiradziev, Kathryn Vanya Ewart

**Affiliations:** Department of Biochemistry & Molecular Biology, Dalhousie University, PO Box 15000, Halifax NS, B3H 4R2, Canada

**Keywords:** antifreeze peptide, antifreeze protein, winter flounder, oligomerisation, amphiphilic, chemical crosslinking, size exclusion chromatography, nucleation, thermal hysteresis

## Abstract

Antifreeze proteins and peptides adhere to ice crystals and inhibit their growth, resulting in freezing point depression. AFP6 is an abundant antifreeze peptide in the blood serum of winter flounder (*Pseudopleuronectes americanus*). This 37-residue peptide forms an amphiphilic α-helix with a hydrophobic ice-binding surface. In this study, prediction of the peptide structure suggested the formation of an antiparallel dimer burying the hydrophobic surfaces of both subunits. The oligomerisation of AFP6 was therefore examined in aqueous solutions using size-exclusion chromatography and chemical crosslinking along with ice-binding evaluation. Crosslinking of non-amidated AFP6, followed by denaturing electrophoresis, revealed bands consistent with monomers and dimers, whereas results for the amidated form and a recombinant mutant suggested additional larger forms. Bands consistent with the presence of monomers and dimers were also predominant following non-amidated AFP6 crosslinking under a variety of solution conditions, with smaller proportions of peptide assembled up to tetrameric sizes; however, the larger oligomers appeared to be absent from AFP6 treated with sodium bicarbonate. Isolated oligomers of non-amidated AFP6 produced ice crystal morphologies and thermal hysteresis comparable to those for the unfractionated peptide suggesting similar interaction with ice. Oligomerisation of AFP6 may allow greater solubility of this amphiphilic α-helical peptide in aqueous solutions.

## 1 │ Introduction

Antifreeze peptides and proteins (AFPs) are structurally diverse molecules produced by a large variety of species ^1^. AFPs rest on the surfaces of ice crystals in a geometrically defined manner, thereby impeding ice growth and lowering the solution freezing point non-colligatively ^1^. In this way, AFPs give rise to distinct ice crystal morphologies and measurable solution effects such as thermal hysteresis ^1,2^. Five structurally distinct types of AFP, defined as binding to ice crystals, have evolved in marine teleost fish species and four of these have been associated with biological roles in the cold ocean survival of the species producing them ^3,4^. The molecular features of AFPs in solution and the structural basis for their association with the ice interface require continuing study in order to understand these biochemical adaptations to freezing environments and to inform the design of new ice-modifying agents.

A substantial percentage of proteins appear to self-assemble into oligomers. Overall, 47% of all protein crystal structures were found to involve homo-oligomers ^5^, with some proportion possibly arising as artefacts of crystallization; conversely, the frequency of self-assembly in yeast (*Saccharomyces cerevisiae*) and human proteomes predicted by modelling was suggested to be approximately 20% ^6^. Furthermore, numerous proteins previously regarded as monomeric were subsequently shown to oligomerise under specific physiological conditions that presumably shift the equilibrium toward association ^5,7^. The quaternary structure of peptides and proteins is foundational to understanding their solution behaviour and mechanisms of action ^6^. For example, oligomerisation can bring structural stability, enhanced binding affinity, and additional mechanisms of regulation ^8^, which may allow self-assembled peptides and proteins greater activity or expansion of their biological roles.

Although the fish AFPs have been largely considered to be monomeric, there are several examples of native oligomers and related structures among them. The smallest antifreeze glycoprotein of Greenland cod (*Gadus ogac*) was found to form dimers and larger aggregates at concentrations above 20 mM, leading to a disproportionate increase in thermal hysteresis activity ^9^. The globular and typically monomeric type III AFP of eelpouts has been proposed to self-assemble into networks following their binding to ice ^10^, although this has not been shown experimentally. However, a natural tandem duplication of the type III AFP from the Antarctic eelpout (*Lycodichthys dearborni*) was shown to have greater thermal hysteresis activity than the single-copy form of the protein ^11^, and further study showed that intermolecular dimerisation can lead to greater thermal hysteresis when the ice-binding sites are aligned ^12^. Engineered monomeric, dimeric, and circular 12-mer forms of the ocean pout AFP showed a modest increase in the ice-absorption rate and thermal hysteresis with dimerisation and a much greater increase in both measures for the 12-mer form, suggesting a clear activity advantage with a few aligned ice-binding sites. The type II AFPs belong to a superfamily of C-type lectins ^13^. Native dimerisation of the smelt (*Osmerus mordax*) AFP was reported, although the possibility of a role in thermal hysteresis activity was not investigated ^14^. The type II AFP of longsnout poacher (*Brachyopsis rostratus*) was modelled as a decamer by overlaying its modelled momomer structure on the decameric structure of a homologous snake venom lectin to explain a remarkable activity increase with concentration ^15^, although the propensity for this AFP to assemble in this manner in solution is unknown.

The small type I AFPs have been intensely investigated, with a 37-residue α-helical AFP6 peptide (also known as HPLC-6, HPLC6, TTTT, and wfl-AFP6) from winter flounder (*Pseudopleuronectes americanus*) employed as a model AFP for structural and functional studies ^16^. AFP6 was proposed to be monomeric based upon a stable CD spectrum with increasing concentration ^17^. In addition, diffusion NMR results and the stability of ^1^H and ^13^C NMR spectra with diminishing temperature for AFP6 were consistent with the presence of a single stable α helix ^18,19^. Nonetheless, the possibility of AFP6 oligomerisation by side-to-side packing on ice was suggested to enhance the effectiveness of ice binding ^20^. Furthermore, analysis of AFP6 using the SEQSEE package ^21^ predicted the formation of multimers due to the preponderance of hydrophobic residues ^22^. Interestingly, analytical centrifugation of the type I AFP of barfin plaice (*Liopsetta pinnifasciata*) suggested the formation of tetramers and trace levels of an octameric form, which were proposed to confer higher thermal hysteresis activity ^23^. This combination of results and predictions made clear the need for further study.

The current investigation was undertaken to test the hypothesis that AFP6 oligomerises under physiological conditions. Predictions of dimerisation were undertaken for AFP6, which was scored alongside other representative proteins using a software package, the native size of the peptide was evaluated by size-exclusion chromatography, the formation of oligomers was examined using covalent crosslinking followed by electrophoresis, and activities of the larger forms were analysed. The results suggested the formation of dimers, with the emergence of smaller proportions of tetramers under some conditions. Remarkably, the predicted structure and the activity of the dimeric form suggested a physiological role in peptide solubility rather than in adhesion to ice.

## 2 │ Materials and Methods

### 2.1 **│ Prediction of peptide dimerisation using a software package**

Dimerisation predictions were conducted for AFP6 and other sequences using the ColabFold v1.6.1 software package ^24^. The AFP6 sequence was the mature sequence encoded from D45 to R81 in the precursor protein sequence ^25^. The sequences of barfin plaice AFP ^23^, monomeric bovine calmodulin ^26^ and dimeric human galectin-1 ^27^ were included for comparison. Sequences were entered as tandem duplicates with a colon (:) between them to allow modelling as dimers. The relaxation number was set to 5, the template mode was set to PDB100, and all other parameters were in their default settings. The Protein Data Bank (PBD) file generated in the best-scoring AFP6 prediction was rendered in iCn3D ^28,29^ in order to visualise the predicted arrangement of the peptide subunits relative to one another along with their binding interface.

### 2.2 │Reagents and antifreeze peptides

All analyses were conducted at room temperature except where otherwise indicated. Deionised water was used in all procedures and chemicals used were reagent grade.

Recombinant AFP6 (hereafter AFP6) and a mutant form (MutAFP6) were expressed as fusion proteins in *Escherichia coli,* cleaved, and purified as previously described ^30^. The expression strategy used to produce recombinant AFP6 was reported by Chang et al. ^30^ and was based on the method employed for the same peptide by Patel and Graether ^31^, with modifications. Briefly, the fusion protein included a vector-encoded N-terminal 6xHis sequence followed by the yeast (*S. cerevisiae*) small ubiquitin-like modifier (SUMO) and the mature AFP6 sequence, which were encoded by DNA that was synthesised and cloned into a pET-15b plasmid by Bio-Basic Inc. Purification steps on a nickel column and cleavage using SUMO protease were conducted to obtain pure AFP6 as previously delineated ^30^. A second expression construct was designed with four amino acid substitutions in the AFP6-encoding portion (T2S, A17L, T24S, T35S); the resulting pure peptide (Mut-AFP6) had the same isoelectric point and molecular mass as AFP6 but showed no thermal hysteresis activity ^30^. Synthetic amidated AFP6 (AFP6-N) was obtained at a purity of 95% from Biomatik and synthetic non-amidated AFP6, identical to the recombinant peptide and therefore also designated as AFP6, was obtained at the same purity from Synpeptide. Peptide concentrations were calculated from absorbance values at 215 and 225 nm as previously described ^32^.

### 2.3 │ Size exclusion chromatography of AFP6

Synthetic AFP6 was resolved on a 1 x 50-cm column (Econo-Column, Bio-Rad) packed with Sephacryl S100-HR (Cytiva) in phosphate-buffered saline with additional NaCl (PBSS, 10 mM phosphate buffer, pH 7.4 with 2.7 mM KCl and 337 mM NaCl) to reflect the NaCl concentration in flounder plasma during winter ^33^. Fractions were collected using a Gilson FC-80K fraction collector and absorbances were measured on a microspectrophotometer (Denovix). Synthetic AFP6 (2 mg) was prepared in 0.5 mL of PBSS. Ferritin was used as a void volume marker and size standards included bovine serum albumin, carbonic anhydrase, cytochrome c, aprotinin, and cyanocobalamin (vitamin B12) in 0.5 mL PBSS. Elution fractions were calculated for AFP6 and for each standard from the centroids of absorbance values at 230 nm for fractions corresponding to each peak using Microsoft Excel. To calibrate the column, the elution fraction (V_e_) of each standard protein was calculated as a multiple of the void volume (V_o_), which was determined using ferritin. The equation of the line of best fit for the standards was determined and it was then used to calculate the molecular weight of AFP6 based upon its elution fraction. Excel (Microsoft) was used for all calculations.

### 2.4 │ Chemical crosslinking treatment of peptides and analysis of products

#### 2.4.1 │ Crosslinking analysis

Glutaraldehyde was added to preparations of AFP6, MutAFP6, and AFP6-N in buffer resulting in a final peptide concentration of 1.5 mM in diluted PBS (8 mM phosphate buffer, pH 7.4 with 69 mM NaCl and 14 mM KCl). Reactions were incubated for 1 hour and quenched by adding glycine to 200 mM. Controls were treated identically except that an equal volume of water was added instead of glutaraldehyde. In experiments to which additional solutes were added, the final buffer was 15 mM phosphate, pH 7.4, and the final additive concentration was 250 mM. This pH was chosen to reflect normal physiological conditions and to avoid both the polymerisation of glutaraldehyde that is possible at higher pH values and the risk of instability of the Schiff base intermediate in the glutaraldehyde-peptide reaction at lower pH values ^34^, so that direct peptide- glutaraldehyde-peptide complexes would form if oligomerisation were to occur. Following this procedure, peptides were resolved by tricine SDS-PAGE as previously described ^32^. Since AFP6 and related peptides are not stained by Coomassie blue or silver, they were incubated in a curcumin stain and imaged in a FluorChem E system (ProteinSimple) as previously described ^32^. Protein size markers used included the Precision Plus Protein™ Dual Xtra Prestained Standards (Bio-Rad) and the PageRuler^TM^ Unstained Low Range Protein Ladder (ThermoFisher Scientific).

#### 2.4.2 │ Peptide size filtration

To prepare size-filtered samples, microcentrifuge tube filtration units (Vivaspin®, Cytiva) were rinsed with 10 mM ammonium bicarbonate, pH 8.3. The crosslinked peptide was filtered by centrifugation through a 3-kDa molecular weight cutoff filter at 8200 x *g* in a microcentrifuge (Eppendorf 5415 C) to remove glutaraldehyde and glycine and to exchange the buffer to 10 mM ammonium bicarbonate, pH 8.3. The retained peptide was then filtered in the same manner through a 10-kDa molecular weight cutoff to separate AFP6 oligomers from monomers.

#### 2.4.3 │ Evaluation of ice binding activities

Ice crystal morphology and thermal hysteresis were determined using a Clifton Nanolitre Osmometer as previously described ^30^. Data were analysed using Excel (Microsoft) and Prism (GraphPad).

## 3 │ Results and Discussion

Modelling of AFP6 and of other peptides and proteins was conducted in Colabfold ^24^ using parameters that would force the model into a dimer and generate scores for the confidence of the structures and of the interfaces generated. The strategy was intended to compare the reliability of dimer predictions with the expectation that the interfaces of true dimers would be predicted more reliably than those forced upon monomeric proteins, thereby allowing insight into the possibility of AFP6 oligomerisation. The predicted local distance difference test (pLDDT) score is a measure of the per-residue accuracy of the predicted structure ^35^, whereas the predicted template modelling (pTM) score represents the accuracy of the entire structure ^35,36^. The ipTM score is similar to the pTM but predicts the accuracy of the interface between two associated protein chains ^37,38^.

Scores for dimers modelled in ColabFold are shown in Table 1. AFP6 and MutAFP6 showed high pLDDT scores (out of 100) with somewhat lower pTM and ipTM scores (out of 1), suggesting a confident prediction of each AFP helix at the local residue level but more moderate confidence at the overall structure and interface levels. The barfin plaice type I AFP was analysed for comparison, as its oligomerisation has been shown experimentally ^23^. The pLDDT score for this peptide was just above 50, suggesting lower confidence in the structure. The pTM score was very low and the ipTM score of 0.194 was far below the range where oligomerisation might be considered. Nonetheless, this low-confidence model showed tetramers consistent with experimental findings ^23^. It is unclear how to interpret these scores in light of the proposed oligomerisation of the barfin place peptide. Since the barfin plaice AFP appears to be predominantly monomeric with a small proportion in tetrameric form ^23^, the model could reflect a relatively weak association with an equilibrium favouring the monomer. Alternatively, the low ipTM score might extend from the low confidence of the structural prediction for this peptide, both at the residue and overall fold levels. Human galectin-1 was included for comparison as a non-covalent dimer ^27^. The pLDDT scores for this protein were above 90 and the ipTM scores were above 0.9, consistent with a highly confident prediction of the structure and of the dimeric interface. In contrast, the scores for bovine calmodulin, which is monomeric ^26^, showed a relatively confident pLDDT prediction, but with an ipTM below 0.1. This reflected the very poor confidence of a prediction made for a dimer that does not exist. Considered together, the results for these known dimeric and monomeric proteins suggest that the parameters used were appropriate for the prediction of oligomerisation. Therefore, the moderate score for AFP6 suggested reasonable confidence in oligomer formation. The Colabfold predicted structure for AFP6 is shown in Figure 1. The two amphiphilic helices are located side by side and oriented in opposite directions to form a symmetrical dimer. The association interface comprises the hydrophobic face of each helix, which was identified as the ice-binding surface ^39^. Thus, this dimer does not appear to be the active form of AFP6. In contrast to the oligomers of AFPs that increase their thermal hysteresis activities by lining up two or more ice-binding surfaces ^9,11,23,40^, the predicted AFP6 dimer buries them.

**FIGURE 1.**
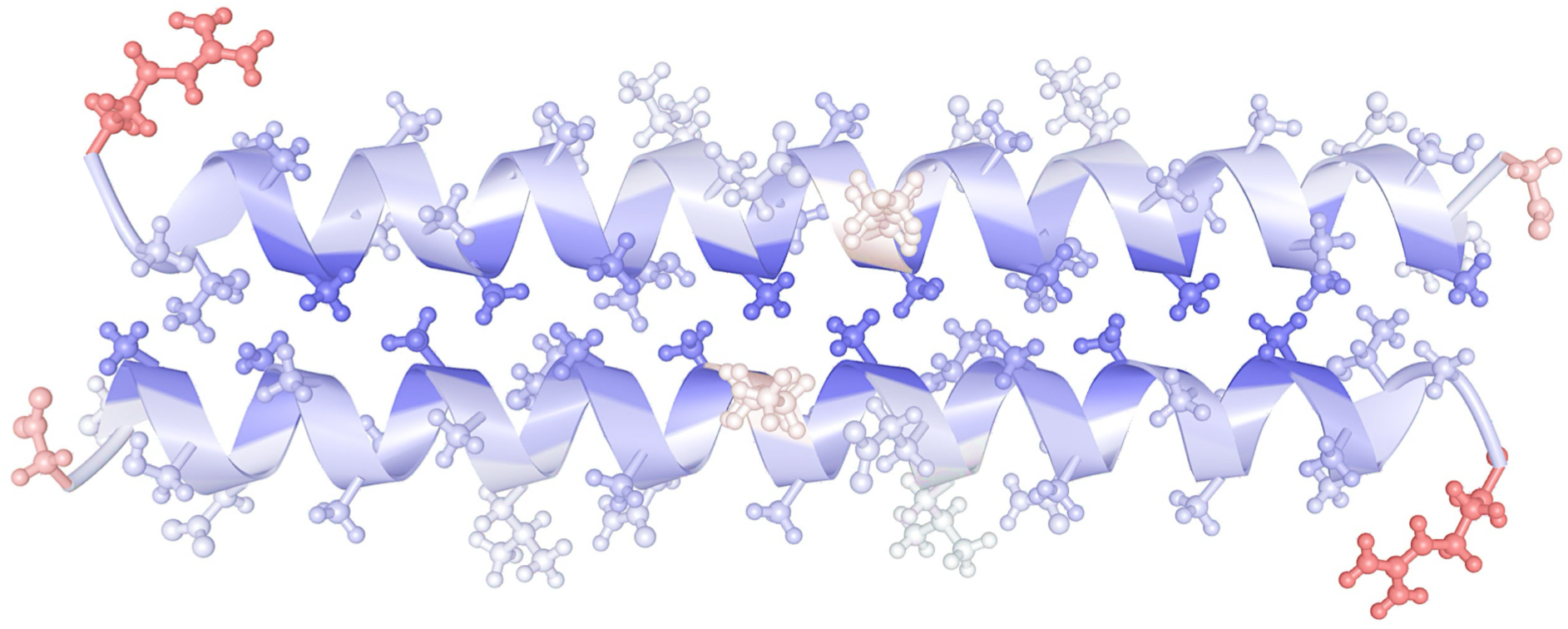
│. Structure of a predicted AFP6 dimer. The dimer was predicted using ColabFold as described in the Methods. The resulting PDB file was visualised as a ribbon diagram with residue side chains shown in ball-and-stick form using iCn3D ^28,29^. Shading represents predicted exposure to solvent with red as 100% and blue as 0%.

**TABLE 1.** │. Predictions of peptide and protein oligomerisation using ColabFold. Parameters used in ColabFold ^27^ were described in the Materials and Methods and were identical for all proteins evaluated. The protein name is given with the acronym used in this report, if applicable. The pLDDT is the predicated local distance difference test score, the pTM is the predicted template modelling score and the ipTM is the interface predicted template modelling score ^24,37^. Data for the models with the highest pLDDT scores for each protein are shown here. For AFP6, the mature protein sequence used extended from D45 to R81 in the protein sequence at Q03659 in the UniProt database, as the pre and pro sequences and the C-terminal amidation were not included.

| Protein name, abbreviation where applicable, and reference | <u>Uniprot sequence</u><br>accession number* | <u>pLDDT</u> | <u>pTM</u> | <u>ipTM</u> |
| --- | --- | --- | --- | --- |
| Antifreeze protein 6 (AFP6), winter flounder <sup>25</sup> | Q03659 | 88.8 | 0.729 | 0.646 |
| Mutated AFP6 (MutAFP6) <sup>30</sup> |  | 85.3 | 0.693 | 0.570 |
| AFP (type I), <u>barfin plaice</u> <sup>23</sup> |  | 52.0 | 0.363 | 0.194 |
| Galectin-1, human <sup>27</sup> | P09382 | 97.4 | 0.930 | 0.910 |
| Calmodulin, bovine <sup>26</sup> | P62157 | 72.4 | 0.331 | 0.074 |

\* Numbers are provided where available.

The possibility of AFP6 dimerisation *in vitro* was first examined by size-exclusion chromatography. A representative chromatogram is shown in Figure 2A. Native AFP6 eluted from the column in a single absorbance peak measured at 230 nm with absorbance values below the baseline at 280 nm, consistent with the absence of aromatic residues in the peptide. The calibration curve obtained for the column using a series of standards, overlaid with elution positions for two AFP6 chromatograms, is shown in Figure 2B. The average of the two molecular weight values calculated for AFP6 from the standard curve was 13.0 kDa. This was similar to previous results of 10 and 12 kDa obtained from size exclusion chromatography of mixtures of AFP6 and related peptides conducted under denaturing conditions ^41,42^. The discrepancy between the molecular weight of 3243 Da, based upon the sequence of AFP6, and the larger one determined by chromatography can be expected in native size-exclusion chromatography because the peptide is a linear α helix ^43^. The extended rod-like structure of AFP6 would result in a greater hydrodynamic radius when compared to more compact or globular proteins. The effect of the extended structure of a peptide on its hydrodynamic radius can be estimated by comparing it with the hydrodynamic radii of 15 monomeric proteins in their native forms in the absence of urea and in their extended, denatured forms in 8 M urea ^44^. The denatured proteins had an average radius 87% greater than that of their native counterparts ^44^. Applying this relationship to AFP6 in order to account for its extended structure, the corrected molecular weight of monomeric AFP6 would be predicted to be 6.1 kDa. Therefore, the extended form of AFP6 cannot account for its elution corresponding to a 13.0-kDa size, which is slightly more than double the corrected molecular weight, suggesting that the peptide may also be forming larger structures. If the peptide forms a side-by-side dimer as shown in Figure 1, the length would be unchanged; however, the greater lateral dimensions of such a dimer might alter the hydrodynamic radius in other ways that affect its chromatographic behaviour.

**FIGURE 2.**
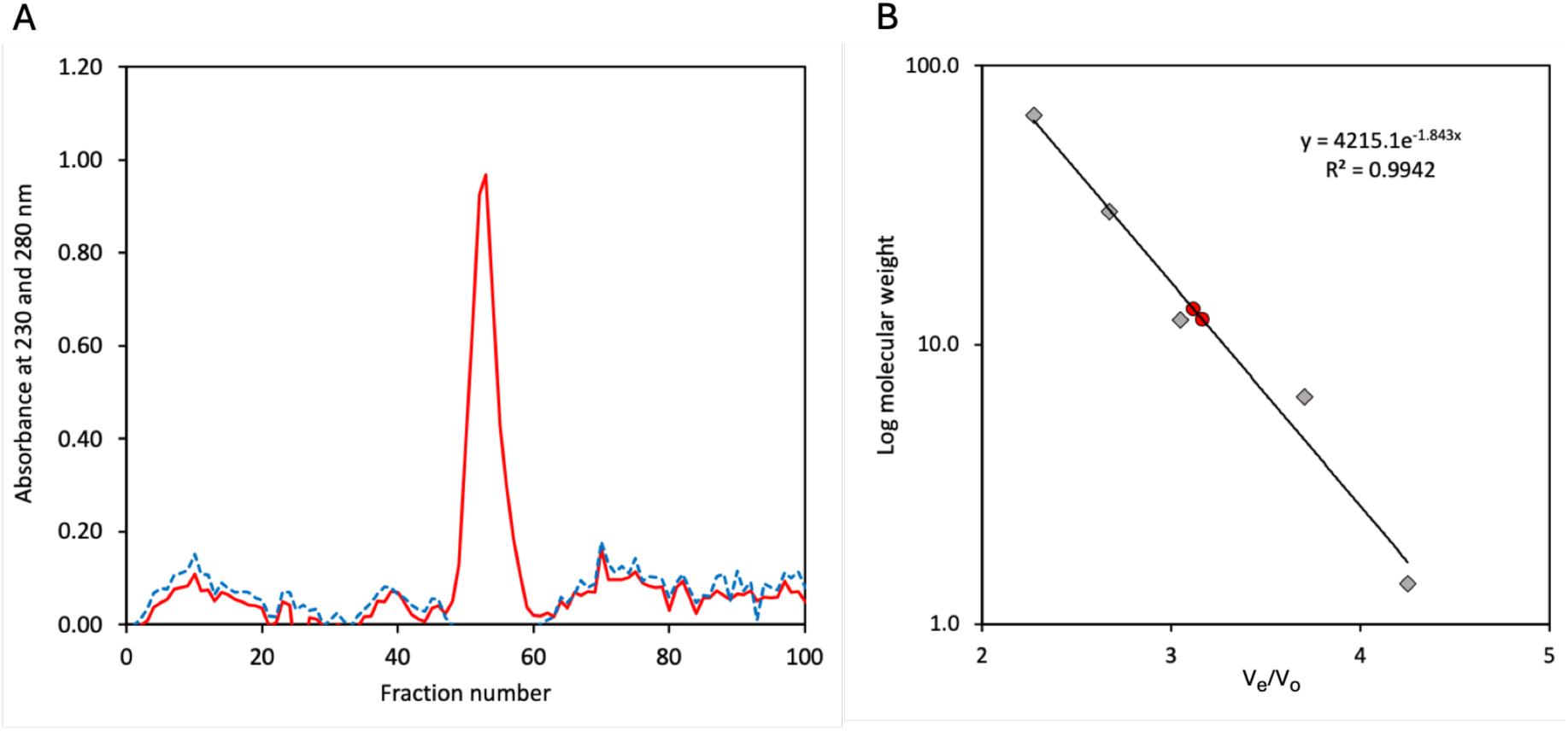
│. Size exclusion chromatography of AFP6. Synthetic AFP6 was applied to a high-resolution Sephacryl S-100 HR column under normal pressure. Panel A shows a representative chromatogram for AFP6. The absorbance at 230 nm is shown as a solid red line and the absorbance at 280 nm is shown as a hatched blue line. Values below 0 are not included. Panel B shows the calibration curve based on the elution fractions calculated for each standard as a multiple of the elution fraction of ferritin (void). V_e_/V_o_ on the horizontal axis represents the elution fraction of each standard (V_e_) as a multiple of the void (ferritin) fraction (V_o_) and the log of the molecular weight is on the vertical axis. Values for standards including bovine serum albumin, carbonic anhydrase, cytochrome c, aprotinin, and cyanocobalamin (in order from smallest to largest V_e_/V_o_ value), which were used to determine the line of best fit, are shown in black circles. The equation of the line of best fit is shown along with the R^2^ value. The V_e_/V_o_ values for AFP6 in two chromatography runs are shown in red circles, which are shown directly on the line of best fit because this line was used to determine the molecular weight in each case.

Covalent crosslinking experiments were conducted on AFP6, AFP6-N, and MutAFP6 to allow any oligomeric forms to retain their native association under denaturing conditions. Peptides treated in this way can be examined for oligomerisation by evaluating their migration on tricine SDS-PAGE. Glutaraldehyde-treated AFP6, AFP6-N and MutAFP6 all showed slower migrating bands than the untreated controls on the resulting stained gels (Figure 3A,B). AFP6 formed dimers, whereas AFP6-N formed primarily dimers with a small proportion of larger structures consistent with trimers and tetramers (Figure 3B). Glutaraldehyde-treated MutAFP also showed larger assemblies (Figure 3B). Nonetheless, each peptide showed a portion remaining in monomeric form. This could not be attributed to limiting amounts of glutaraldehyde, as its highest concentration (1.5%) was approximately 50 to 100-fold greater than those of the peptides on a molar basis. Instead, it suggests an equilibrium between monomeric and dimeric or more complex forms that may be poised to shift in response to peptide concentrations or solution conditions. The concentrations of peptide used were well below the 3–5 mM accumulation of AFP present in winter flounder plasma during the coldest months ^37,38^ so that any interaction among molecules would reflect their association at physiological concentrations.

**FIGURE 3.**
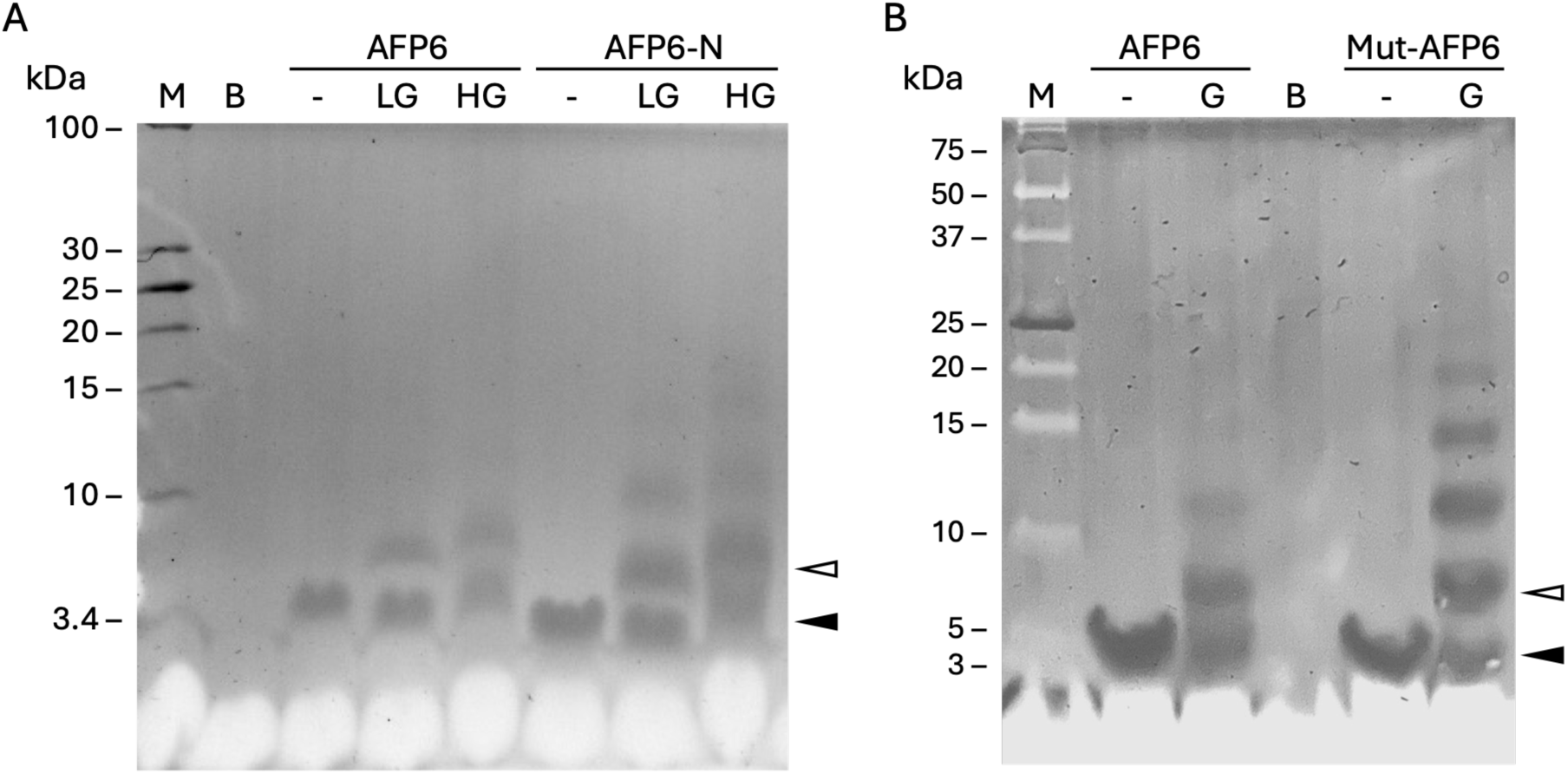
│. Tris-tricine SDS-PAGE analysis of crosslinked antifreeze peptides. Samples were resolved on an 18% acrylamide gel and stained with curcumin. Panel A: The PageRuler^TM^ Unstained Low Range Protein Ladder (ThermoFisher Scientific) is in lane M and buffer is in lane B. For other lanes, AFP6 is the recombinant AFP6, AFP6-N is synthetic amidated AFP6, “-” indicates no glutaraldehyde, LG indicates 0.15% glutaraldehyde, and HG indicates 1.5% glutaraldehyde. Panel B: The Precision Plus Protein™ Dual Xtra Prestained Standards (Bio-Rad) are in lane M. For other lanes, AFP6 is recombinant AFP6, Mut-AFP6 is the mutant recombinant AFP6, “-” indicates no glutaraldehyde and G indicates 1.5% glutaraldehyde. In both panels, the solid and open arrows show the positions of monomeric and dimeric AFP6 peptide, respectively.

The oligomerisation of AFP6 was investigated for peptide variants and under multiple conditions. Oligomers similar to those in the control solutions appeared to form in the presence of additional solution components, which included osmolytes, a crowding agent, non-ionic detergent and salts (Figure 4). The control solution for this experiment was 15 mM phosphate buffer, pH 7.4, which allowed the role of salt in the behaviour of AFP6 in flounder blood plasma to be tested. Although the concentration of added NaCl was 250 mM in this experiment, dimers remained the dominant oligomeric form with only a slight increase in the proportion of structures larger than dimers (Figure 4). Sodium bicarbonate at the same concentration appeared to diminish the formation of oligomers larger than dimers. The non-ionic detergent Triton X-100 had no discernible effect on AFP6 oligomerisation, although it interfered with the resolution of the monomeric peptide in the gel (Figure 4). The effect of temperature on oligomer formation by AFP6 revealed monomers and dimers to be the major forms of AFP6 at 0 and 4 °C (Figure 5), supporting the notion that AFP6 assembles into dimers under natural winter conditions rather than forming larger structures. Although the reaction of glutaraldehyde with protein functional groups is slower at lower temperatures, the 1-hour reaction time used in these experiments should allow all stable oligomers to be crosslinked, as far shorter incubation invervals have been used successfully for this reaction ^45,46^. An unidentified band migrating just ahead of the 25-kDa marker was present in both the glutaraldehyde-treated and untreated samples at higher temperatures, but it did not interfere with interpretation of the AFP6 results (Figure 5). Furthermore, additional bands too large to resolve in the separating gel were visible at the top of the gel in the cross-linked samples at 20 and 40°C (Figure 5). These bands, which were consistent with the presence of large aggregates, were not evident in the peptide without glutaraldehyde or in the peptide incubated in the presence or absence of glutaraldehyde at lower temperatures (Figure 5). Thus, these unusual bands may represent denatured AFP6 aggregates that are crosslinked at higher temperatures. The melting point of AFP6 was found to be 31 °C ^32^; however, with a relatively broad melting curve ^17,32^, a minor portion of AFP6 appears to be unfolded at 20°C while a small proportion appears to retain helical structure at 40 °C ^32^. Unfolded peptide might form non-native aggregates, which could then be cross-linked by glutaraldehyde and potentially form large covalent structures. However, if present, this denatured material would not be representative of the native assembly of AFP6. Considered together, these results suggest that AFP6 can form dimers in solutions at temperatures and salt concentrations that correspond to those in flounder blood during winter ^33,47^.

**FIGURE 4.**
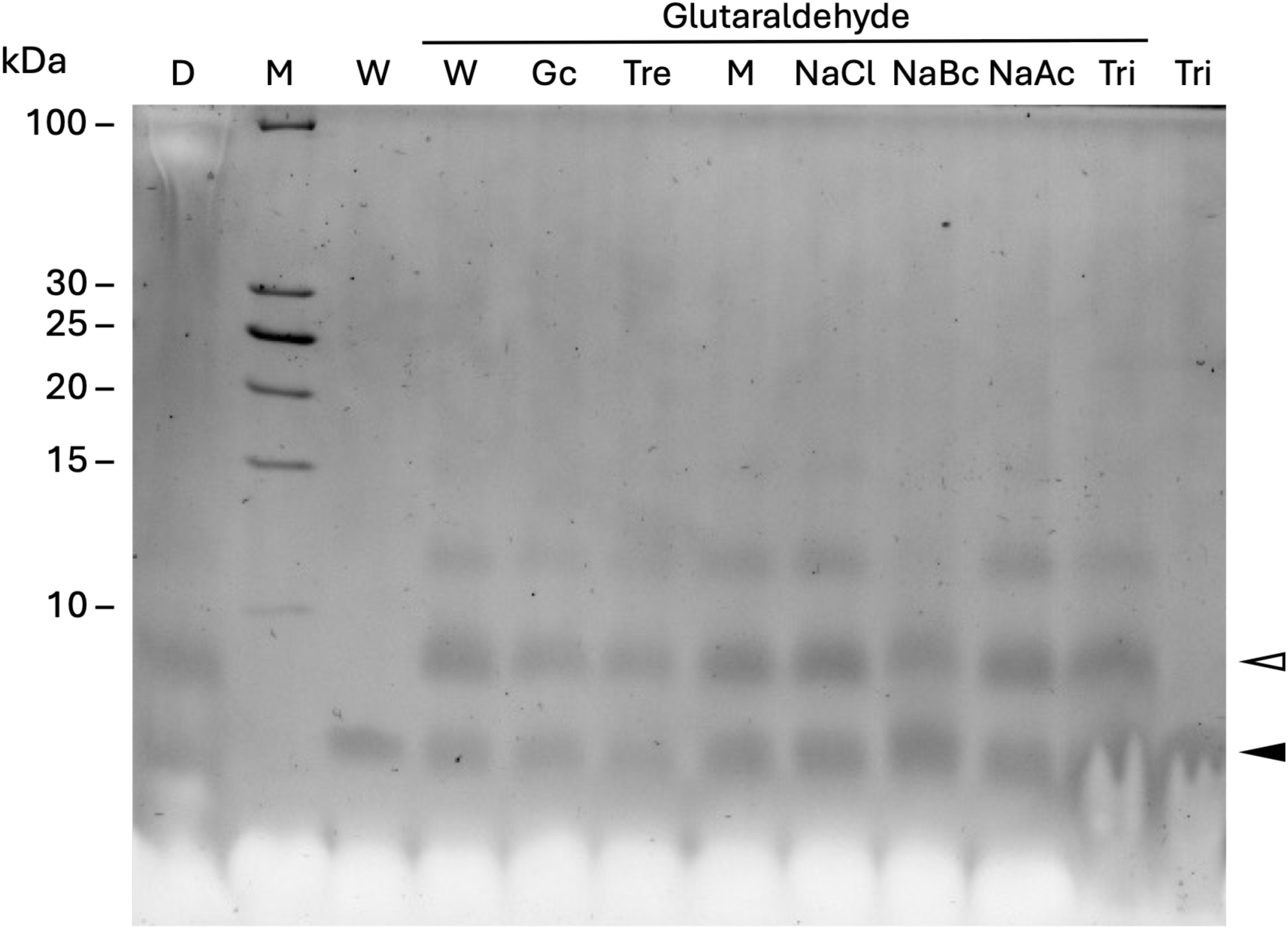
│. Tris-Tricine SDS-PAGE analysis of recombinant AFP6 crosslinked in the presence of added solution components. Samples were resolved on an 18% acrylamide gel and stained with curcumin. The PageRuler^TM^ Unstained Low Range Protein Ladder (ThermoFisher Scientific) is in lane M. Other lanes show recombinant AFP6 in 15 mM phosphate buffer, pH 7.4 with 250 mM of the following components where indicated: dextran sulphate (D), glycerol (Gc), trehalose (Tre), mannitol (M), sodium chloride (NaCl), sodium bicarbonate (NaBc), sodium acetate (NaAc), Triton X-100 (Tri), and the controls to which water was added (W). Glutaraldehyde-treated samples are indicated. The solid and open arrows show the position of monomeric and dimeric peptide, respectively.

**FIGURE 5.**
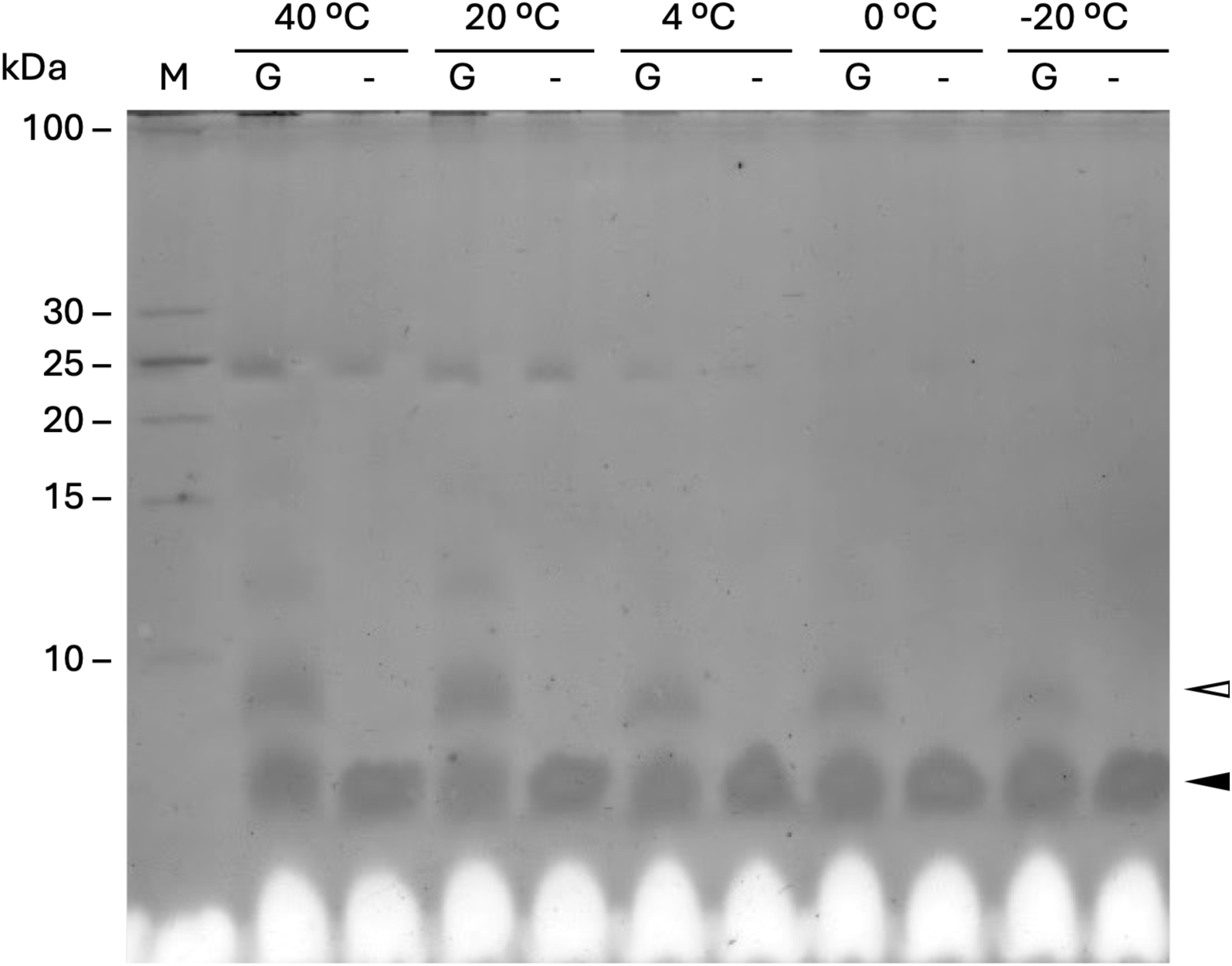
│. Tris-Tricine SDS-PAGE analysis of recombinant AFP6 crosslinked at different temperatures. M is the PageRuler^TM^ Unstained Low Range Protein Ladder (ThermoFisher Scientific). G indicates 1.5% glutaraldehyde and “-” indicates no glutaraldehyde. Incubation temperatures are indicated. The solid and open arrows show the position of monomeric and dimeric peptide, respectively.

The antifreeze activity of the cross-linked oligomeric form of AFP6 was evaluated. Cross- linked AFP6 retained on a 10-kDa filter was shown to consist primarily of dimers with no evidence of monomer presence, whereas the portion that passed through the filter consisted of monomeric peptide with only a trace amount of the oligomeric form (Figure 6A). The ice binding activity of the crosslinked oligomeric AFP6 was similar to that of the untreated AFP6, both in terms of the thermal hysteresis and ice crystal morphologies generated by these peptides (Figure 6B,C). The crosslinked dimeric AFP6 could have been expected to have impaired thermal hysteresis activity since the ice-binding surfaces are predicted to be buried in the dimer (Figure 1). However, the flexibility of the crosslinked dimer is unclear and may depend on the positions of glutaraldehyde linkages. It is possible, for example, that AFP6 dimers covalently linked by glutaraldehyde at Lys18 could still freely dissociate, thereby allowing one or both ice-binding faces to make contact with ice, while remaining tethered to one another.

**FIGURE 6.**
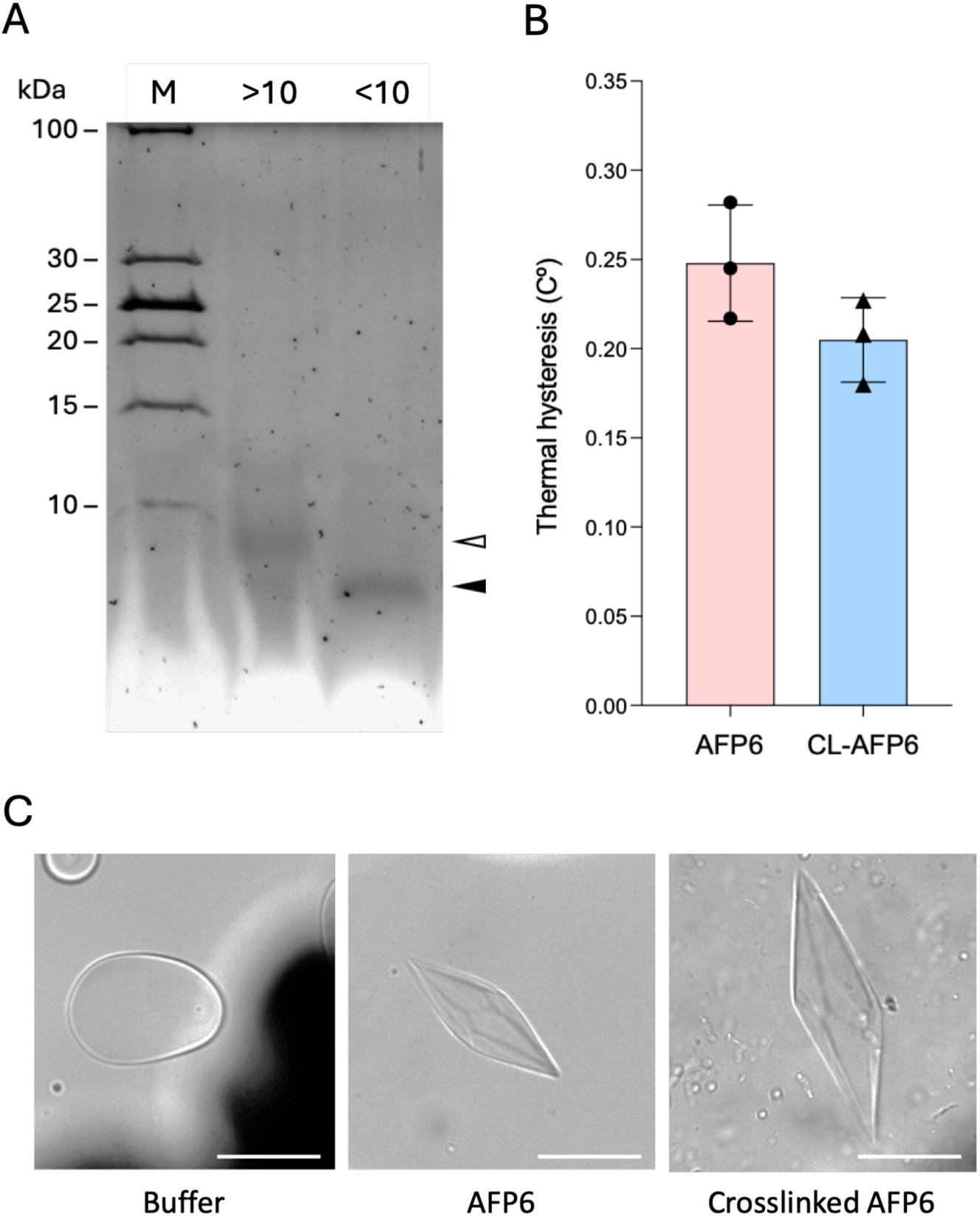
│. The effect of dimer crosslinking on antifreeze activity of recombinant AFP6. In Panel A, M is the PageRuler^TM^ Unstained Low Range Protein Ladder (ThermoFisher Scientific), “<10” is the 10-kDa filter filtrate containing monomers, and “>10” is the 10-kDa filter retentate containing mainly dimers. The solid and open arrows show the position of monomeric and dimeric peptide, respectively. In Panel B, thermal hysteresis of untreated AFP6 and crosslinked dimeric AFP6 are shown. Data shown are the mean ± SD for triplicate readings with symbols representing individual values. In Panel C, representative ice crystal morphologies of ice grown in the presence of untreated AFP6 and crosslinked dimers are shown. White bars in the images represent 50 μm.

It is interesting that oligomerisation of AFP6 was predicted by modelling and observed by crosslinking in this study when previous biophysical investigations suggested the peptide to be monomeric. These differences may be due to the contrasting conditions and methods used. The consistency in the CD spectrum with increasing concentration of AFP6 was interpreted as an indication of monomeric character ^17^; however, it may reflect the presence of native α helices in the AFP6 dimer structure predicted using ColabFold (Figure 1). The CD spectrum, typical of an α helix, could therefore be expected to remain stable over the course of AFP6 dimer assembly. In addition, the maximal concentration used in the CD experiment ^17^ was approximately 10-fold lower than the physiological concentrations used in the current study and may therefore have been insufficient for dimerisation to occur. Diffusion NMR studies employing a typical concentration (4.8 mM) of AFP6 also suggested a monomeric peptide. However, in contrast to dimers formed by globular subunits, which should result in substantial changes in the diffusion coefficient, the dimer predicted for AFP6 composed of antiparallel adjacent α helices with a buried surface (Figure 1) might not give diffusion results readily distinguishable from those for a monomeric helix. Furthermore, AFP6 was interpreted to be monomeric based upon 13C- and 1H-NMR analyses conducted on peptide dissolved in pure water ^18^. In the absence of pH buffering and ions to screen electrostatic repulsion, the monomeric form might be favoured in water even if a peptide had a propensity for oligomerisation in physiological solutions. Therefore, the contrast in peptide assembly among these studies could reflect differences among the parameters measured, the concentrations of peptide, and the solution conditions.

Although the predicted dimer structure for AFP6 precludes ice binding (Figure 1), the dimerisation of AFP6 could serve a distinct purpose in support of the freezing point depression that is required by the fish to survive. The moderately active AFPs of fish must be present at high concentrations, relative to most other proteins in the blood plasma or in tissues, in order to confer meaningful freezing point depression ^4,13^. As noted above, in the winter flounder, AFP6 can attain concentrations of 3–5 mM ^13^. In order to reach these levels, many species have evolved multigene families encoding the AFPs they require for survival ^4^. Therefore, one component of an AFP adaptation is the accumulation of sufficient protein. Yet, in this context, solubility could become limiting. The apolipoprotein-like type IV AFP of longhorn and shorthorn sculpins (*Myoxocephalus octodecemspinosus* and *M. scorpius*, respectively) ^3,48^ may be an example of this limitation. The type IV AFPs show activity *in vitro* but do not reach the concentrations needed to bring about protection from freezing in the sculpins, possibly because of low protein solubility ^3^. In contrast, AFP6 is a highly abundant AFP isoform present in flounder blood along with several similar isoforms ^49^. The hydrophobic surface of AFP6 that binds ice could also limit the solubility of the monomer; however, dimerisation of AFP6 could enhance the solubility of this peptide by masking its hydrophobic ice-binding face and thereby allowing greater concentrations to be attained. In the analogous situation of the amphiphilic α-helical antimicrobial peptides, as the non-polar face becomes more hydrophobic and the monomer solubility decreases, dimerisation has been suggested to increase ^50^. Still, the formation of AFP6 dimers predicted to bury their hydrophobic ice-binding surfaces raises the question of how the monomeric peptide might become available to interact with ice. Crosslinking results suggest that a substantial proportion of AFP6 remained monomeric in solution under all conditions tested (Figures 2–5); however, in the ice-water interface layer, only monomeric AFP6 can interact with ice ^51^. This process would sequester monomers, thereby shifting the local monomer-dimer equilibrium in the ice-water interface. In this way, dimers that are present in solution could give way to monomers compatible with ice-binding precisely where they are needed.

## Acknowledgments

We thank Aja Deeble, Jenny Ho, and Zulqi Memon for technical assistance. We are grateful to Andrew Roger for the use of his gel imager, to Jamie Kramer for access to software for editing, and to many people in the Dalhousie Faculty of Medicine for donations of excess dry ice. We also thank H. Stephen Ewart for proofreading and assistance with formatting of the manuscript.

## Author contributions

Conception and project design: LLV and KVE; experimental design and data acquisition: LLV, AO, XJC, JS, ML, AH, BK, and KVE; data analysis and interpretation of results: LLV, AO, JS, ML, AH, BK, and KVE; Original manuscript draft preparation: LLV and KVE. All authors reviewed the results and approved the manuscript.

## Data availability

All original data can be made available by the corresponding author upon reasonable request.

## Abbreviations

AFP, antifreeze protein or antifreeze peptide; NMR, nuclear magnetic resonance; SEC, size exclusion chromatography; SDS-PAGE, sodium dodecyl sulphate- polyacrylamide gel electrophoresis.

